# Posttranscriptional regulation of intestinal epithelial cell repair by RNA binding protein IMP1

**DOI:** 10.1101/368050

**Authors:** Priya Chatterji, Kelly A. Whelan, Sarah F. Andres, Fernando C. Samper, Lauren A. Simon, Rei Mizuno, Emma T. Lundsmith, David S.M. Lee, Shun Liang, H.R. Sagara Wijeratne, Stefanie Marti, Lillian Chau, Patrick A. Williams, Veronique Giroux, Benjamin J. Wilkins, Gary D. Wu, Premal Shah, Gian G. Tartaglia, Kathryn E. Hamilton

## Abstract

RNA binding proteins, such as IMP1, are emerging as essential regulators of intestinal development and cancer. IMP1 hypomorphic mice exhibit severe intestinal growth defects, yet it’s role in adult intestinal epithelium is unclear. We employed ribosome profiling to test the effect of IMP1 loss on the “translatome” in colon cancer cell lines. In parallel, we evaluated mice with intestinal epithelial-specific *Imp1* deletion (*Imp1^ΔIEC^*) following irradiation or colitis models. Ribosome-profiling revealed translation efficiency changes for multiple pathways important for intestinal homeostasis, including autophagy, in IMP1 knockout cells. We found increased autophagy flux in *Imp1^ΔIEC^* mice, reinforced through *in silico* and biochemical analyses revealing direct binding of IMP1 to autophagy transcripts *MAP1LC3B* and *ATG3*. We found that *Imp1^ΔIEC^* mice exhibit enhanced recovery following irradiation, which is attenuated with genetic deletion of autophagy gene *Atg7*. Finally, we demonstrated that IMP1 is upregulated in Crohn’s disease patients and *Imp1* loss lessened colitis severity in mice. These studies demonstrate that IMP1 acts as a posttranscriptional regulator of gut epithelial repair post-irradiation and colitis, in part through modulation of autophagy.

## Introduction

Intestinal epithelium maintains its integrity through orchestration of self-renewal, proliferation, differentiation, and cell death during homeostasis and in response to stress. The rapidity with which intestinal epithelium must respond to environmental stressors suggests a necessity for multiple layers of gene regulation. RNA binding proteins (RBPs) have emerged as critical regulators of intestinal proliferation and stem cell dynamics (Li, Yousefi et al., 2015, Madison, Jeganathan et al., 2015, Madison, Liu et al., 2013, Tu, Schwitalla et al., 2015, Wang, Li et al., 2015, Yousefi, Li et al., 2016). *IGF2* mRNA-binding protein 1 (IMP1) is a RBP with primary roles in mRNA trafficking, localization, and stability. Target mRNAs of IMP1 (also called IGF2BP1, CRD-BP, ZBP1) include *IGF2, ACTB, MYC, H19, CD44, GLI1 and PTGS2* (Leeds, Kren et al., 1997, Lemm & Ross, 2002, Manieri, Drylewicz et al., 2012, Nielsen, Christiansen et al., 1999, Noubissi, Goswami et al., 2009, Ross, Oleynikov et al., 1997, Runge, Nielsen et al., 2000, Vikesaa, Hansen et al., 2006). *In vitro* studies demonstrate that IMP1 forms stable complexes with its target mRNAs, confining these transcripts to ribonucleoprotein particles (RNPs) and stabilizing mRNA or inhibiting translation (Bernstein, Herrick et al., 1992, Elcheva, Goswami et al., 2009, Gu, Wells et al., 2008, Huttelmaier, Zenklusen et al., 2005, Noubissi, Elcheva et al., 2006, Noubissi et al., 2009, Stohr, Kohn et al., 2012, Stohr, Lederer et al., 2006, Vikesaa et al., 2006, Weidensdorfer, Stohr et al., 2009). IMP1 also plays a functional role in mRNA transport to aid in various cellular processes, including movement and polarity (Gu, Katz et al., 2012, Vikesaa et al., 2006). Finally, PAR-CLIP and eCLIP studies have identified a myriad of IMP1 targets, providing important insight into the diverse and context-specific roles of IMP1 via regulation of specific transcripts or transcript groups (Conway, Van Nostrand et al., 2016, Hafner, Landthaler et al., 2010).

In mice, *Imp1* is expressed in the small intestine and colon during embryonic development through postnatal day 12 and at low levels during adulthood (Hansen, Hammer et al., 2004). *Imp1* hypomorphic mice exhibit dwarfism, intestinal defects and perinatal lethality (Fakhraldeen, Clark et al., 2015, Hansen et al., 2004). Recent studies in the fetal brain implicate Imp1 as a regulator of differentiation of stem/progenitor cells, where *Imp1* deletion leads to neural stem cell depletion (Nishino, Kim et al., 2013). In adult mouse colon, IMP1 is expressed in the epithelial crypt base and in mesenchymal cells following injury (Dimitriadis, Trangas et al., 2007, Manieri et al., 2012). Our prior published studies demonstrated that IMP1 may promote or suppress colon tumorigenesis based upon its expression and function in the epithelial or mesenchymal compartments, underscoring the notion that IMP1 may exhibit opposing effects in different contexts (Hamilton, Chatterji et al., 2015, Hamilton, Noubissi et al., 2013). Taken together, *in vivo* studies suggest that IMP1 is a key regulator of development and cancer, potentially via regulation of stem/progenitor cell maintenance (Degrauwe, Suva et al., 2016).

Prior *in vitro* reports have suggested a role for IMP1 in cellular stress response. Studies of the IMP1 chicken orthologue, ZBP1, revealed an essential role in the integrated stress response (ISR) via differential regulation of mRNA fates in non-stressed versus stressed cells (Stohr et al., 2006). Characterization of IMP1 RNP granules *in vitro* revealed enrichment of mRNAs encoding proteins involved in the secretory pathway, ER stress, and ubiquitin-dependent metabolism (Jonson, Vikesaa et al., 2007). Furthermore, evaluation of processing bodies (P-bodies) using fluorescence-activated particle sorting (FAPS) demonstrated enrichment of IMP1 together with translationally repressed mRNAs, suggesting that IMP1 may regulate stabilization or repression of target mRNAs (Hubstenberger, Courel et al., 2017). Despite its considerable importance in normal development and its role in coordinating cellular stress, the specific functional roles of IMP1 in adult tissues have yet to be elucidated *in vivo*. In the present study, we tested the hypothesis that IMP1 functions as a regulator of epithelial response to damage in adult tissues.

## Results

### IMP1 knockout reveals global changes in active translation

*Imp1* hypomorphic mice exhibit intestinal defects and perinatal lethality, yet the contribution of IMP1 to normal gut homeostasis and repair is not known (Fakhraldeen et al., 2015, Hansen et al., 2004). We utilized CRISPR-mediated IMP1 deletion in the SW480 colorectal cancer cell line to evaluate “translatome”-wide effects of IMP1 loss (Fig. 1A; S1A). Following deep sequencing to compare total RNA abundance in SW480 cells with and without IMP1, RNA fragments protected by bound ribosomes were sequenced to define actively translating mRNAs (Data deposited in GEO (GSE112305) [NCBI tracking system #18999297]). We found that *IMP1* loss affected gene expression on both transcriptional and translational levels. Of the 10043 genes analyzed, we saw no change in RNA level or ribosome binding in 7386 genes (Figure 1B). 642 transcripts were exclusively differentially regulated at the transcription level whereas 1264 genes were only differentially regulated at the translational level. Furthermore, in 28 genes, translation direction was completely antagonistic to transcription, whereas translation reinforced transcriptional direction in 60 genes (Figure 1B). Additionally, in 663 genes translation acted as a buffering mechanism (McManus, May et al., 2014), whereby protein levels remain constant despite changes in mRNA levels.

**Figure 1.**
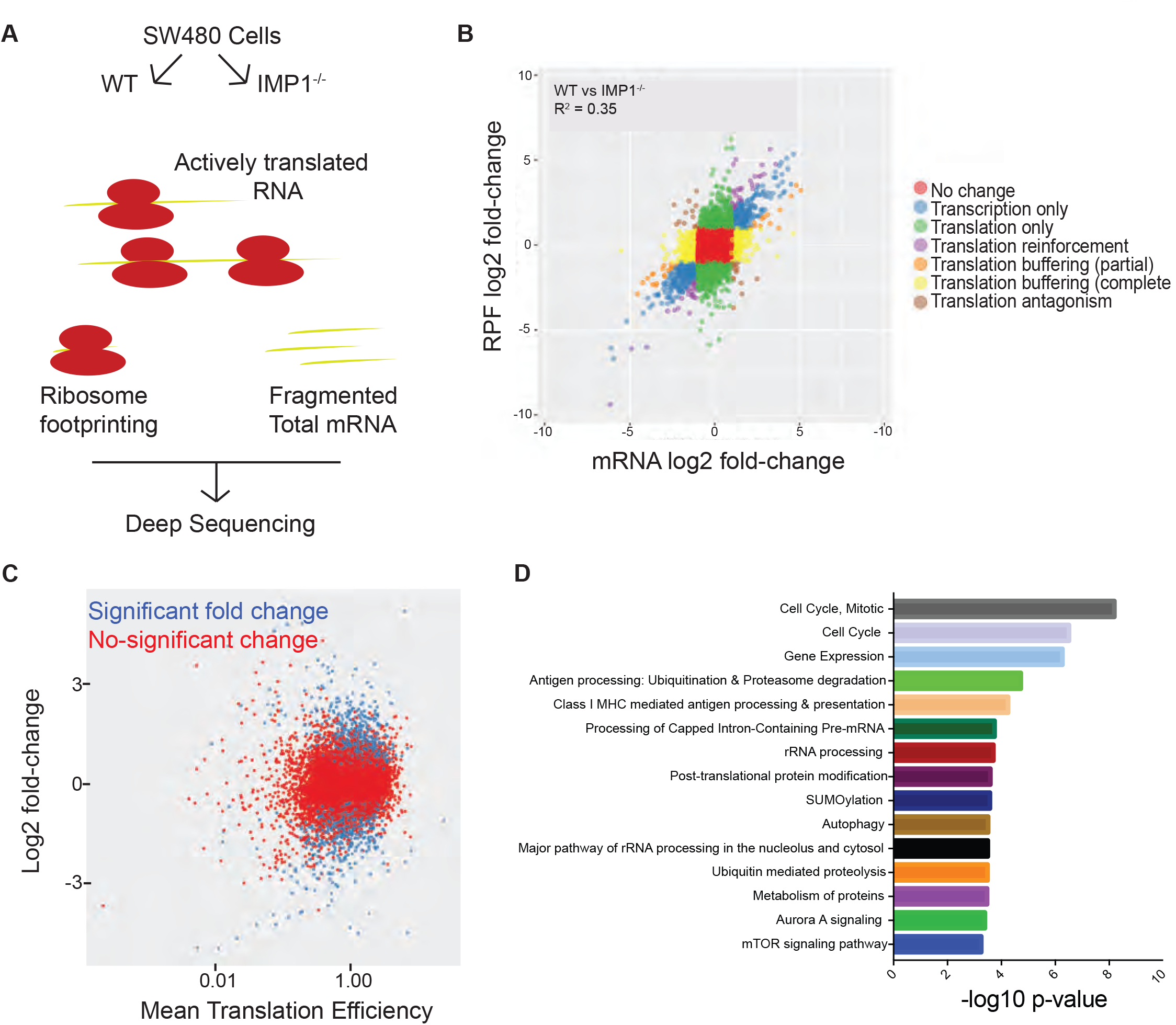
IMP1 knockout reveals global changes in active translation. **A**. Simplistic schematic of ribosome profiling technique. **B**. Scatterplot of differential expression between SW480 cells with and without *IMP1* deletion. The log2 fold change between ribosome-bound RNAs (ribosome protected fragments, or RPF) and total mRNA is plotted. The plot indicates that IMP1 regulates both mRNA abundance and translation. **C**. Scatterplot of genes with significant (in blue) differential translational efficiencies between SW480 cells with and without *IMP1* deletion. Translation efficiencies of transcripts are calculated as the ratio of reads of ribosome-protected fragments to the reads in total mRNA abundance. **D**. Pathway analysis using Toppgene gene enrichment analysis software of differentially expressed genes from **C** to define which signaling/effector pathways are enriched with *IMP1* deletion.

Translational efficiency (TE) of a gene is defined as the ratio of abundance in ribosome protected fragments (RPF) to that of total mRNA abundance for a gene (Zhong, Karaletsos et al., 2017). We compared TE changes between the two genotypes and observed differential TE for 1469 genes (Figure 1C, Table S1). Pathway enrichment analysis for TE genes (Chen, Bardes et al., 2009) differentially expressed in cells with IMP1 deletion revealed a significant representation of pathways linked to cell cycle, gene expression and RNA processing, post-translational modification, autophagy, and metabolism (Figure 1D, Table S2). These analyses support the notion that IMP1 may exhibit pleiotropic effects and that evaluating tissue-and context-specific outcomes permits a deeper understanding of IMP1 functions.

### Mice with Imp1 loss exhibit morphological changes in Paneth cells and enhanced autophagy

Evidence in *Imp1* hypomorphic mice, which express only 20% of wild type Imp1, suggests that its expression is critical for normal growth and development, as these mice exhibit high perinatal mortality and morphological defects in intestine epithelium (Fakhraldeen et al., 2015, Hansen et al., 2004). We therefore evaluated *VillinCre; Imp1^fl/fl^* mice (*Imp1^ΔIEC^* mice), in which IMP1 is deleted solely in intestinal and colonic epithelium (Hamilton et al., 2013). We did not observe gross phenotypic differences in small or large intestine between *Imp1^WT^* (floxed, but intact alleles) and *Imp1^ΔIEC^* mice, suggesting that intestinal epithelial IMP1 is dispensable during homeostasis (Fig. S2A). We analyzed the number and morphology of differentiated epithelial cell types and although there was no difference in total Paneth, goblet, or enteroendocrine cell numbers between genotypes (Fig. S2B, C), we observed diffuse lysozyme staining in *Imp1^ΔIEC^* Paneth cells (Fig. 2A-C). Autophagy gene mutations have been associated with Paneth cell granule defects in Crohn’s disease patients (Liu, Gao et al., 2014, Liu, Gurram et al., 2016, Liu, Naito et al., 2017, Stappenbeck & McGovern, 2017, VanDussen, Liu et al., 2014). Furthermore, we found changes in autophagy pathway TE in ribosome profiling analysis. We therefore evaluated autophagy in *Imp1^ΔIEC^* mice. We performed western blotting for cleaved LC3 in freshly isolated *Imp1^ΔIEC^* colon and jejunum crypt cells. A shift from the upper to lower LC3 band suggested enhanced autophagy flux in *Imp1^ΔIEC^* mice (Fig. 2D; Fig.S2D). We observed concurrently a decrease in autophagy cargo-associated protein p62 in *Imp1^ΔIEC^* colon crypts (Fig. 2D). To confirm changes in autophagy flux, we again utilized freshly isolated, live crypt cells and stained with the cationic amphiphilic tracer dye CytoID, which incorporates into autophagic structures. We validated CytoID as a tool to measure autophagy using *Atg7^ΔIEC^* mice (Fig. S3A, B). CytoID staining and flow cytometry analysis revealed an increase in basal autophagic vesicle content in *Imp1^ΔIEC^* mice compared to *Imp1^WT^*, an effect that was amplified in mice at the day 4 recovery time point following 12Gy irradiation (Fig. 2E). Taken together, these data suggest a modest, yet significant effect of IMP1 deletion to enhance autophagy flux in intestinal crypts.

### IMP1 interacts with autophagy transcripts

To confirm a direct role for IMP1 knockdown to induce autophagy, we evaluated Caco2 cells transfected with IMP1 siRNA and observed a robust increase in LC3-I/LC3-II (Fig. 3A). Thus far, we demonstrated that Imp1 loss: 1) broadly affects the autophagy pathway at the translational level (Fig. 1D), and 2) leads to increased LC3 protein expression (Fig. 2D, S2D, 3A). One way in which RBPs regulate gene expression is via direct binding of target transcripts. We therefore evaluated direct binding of IMP1 to autophagy transcripts. We first performed *in silico* analyses to assess binding propensities of IMP1 for autophagy transcripts using catRAPID, which predicts RNA:protein interactions based upon nucleotide and polypeptide sequences as well as physicochemical properties (Cirillo, Blanco et al., 2016). These analyses predicted binding of IMP1 to *BECN1, MAP1LC3B and ATG3* transcripts (Fig. 3B), as well as positive control *ACTB*. Lower relative binding scores were predicted for *ATG16L1, ATG7, and ATG5*. This algorithm predicted no binding to negative targets *TNFRSF1B and ITGA7*(Conway et al., 2016). We next evaluated published eCLIP data (Conway et al., 2016) in human pluripotent stem cells for the same autophagy transcripts and found that they were bound by IMP1 (Fig. 3C). We also found a signification correlation between catRAPID-predicted binding and eCLIP binding (data not shown).

**Figure 2.**
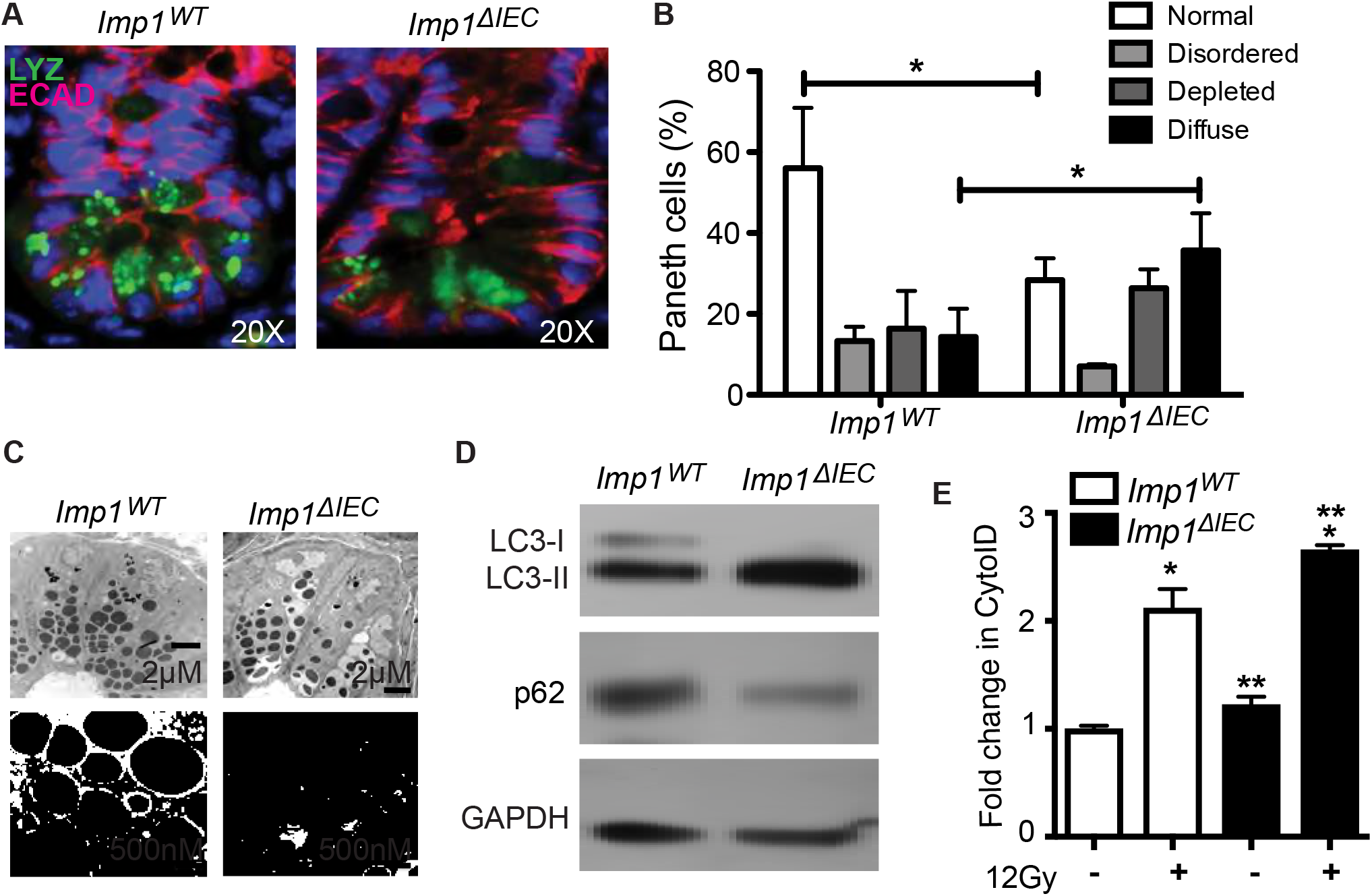
Mice with Imp1 loss exhibit morphological changes in Paneth cells and enhanced autophagy. A. Paneth cells from *Imp1^WT^* and *Imp1^ΔIEC^* mice were evaluated histologically using IF for lysozyme (LYZ). E-cadherin (ECAD) staining was used to demarcate individual epithelial cells. Note presence of diffuse lysozyme staining in *Imp1^ΔIEC^* mice. **B**. Published lysozyme scoring was utilized to evaluate specific Paneth cell phenotypes. *Imp1^ΔIEC^* mice exhibit a significant shift from normal to diffuse lysozyme phenotype (n= 4 mice per genotype). **C**. Transmission electron microscopy revealed an abundance of small, electron dense granules in *Imp1^ΔIEC^* mice. **D**. To confirm a direct role for IMP1 knockdown to induce autophagy, we evaluated epithelial cells from colon in *Imp1^WT^* and *Imp1^ΔIEC^* mice. This confirmed a robust increase in LC3 and decrease in p62, which together indicate enhanced flux. Blots are representative of 3 independent experiments. **E**. Live cell staining of autophagic structures with the cationic amphiphilic tracer dye CytoID indicated a significant increase in autophagic vesicles in crypts from *Imp1^ΔIEC^* mice (n=8) compared to *Imp1^WT^* mice (n= 7) using flow cytometry. There was a modest increase in basal CytoID in *Imp1^ΔIEC^* enteroids compared to controls, as seen with FACS analysis of isolated crypts from *Imp1^ΔIEC^* mice, and both genotypes exhibited an increase in CytoID puncta following 12Gy radiation treatment. (All data are expressed as mean ± SEM. *, p < 0.05; **, p < 0.01; by ordinary one-way ANOVA test or standard t-test).

**Figure 3.**
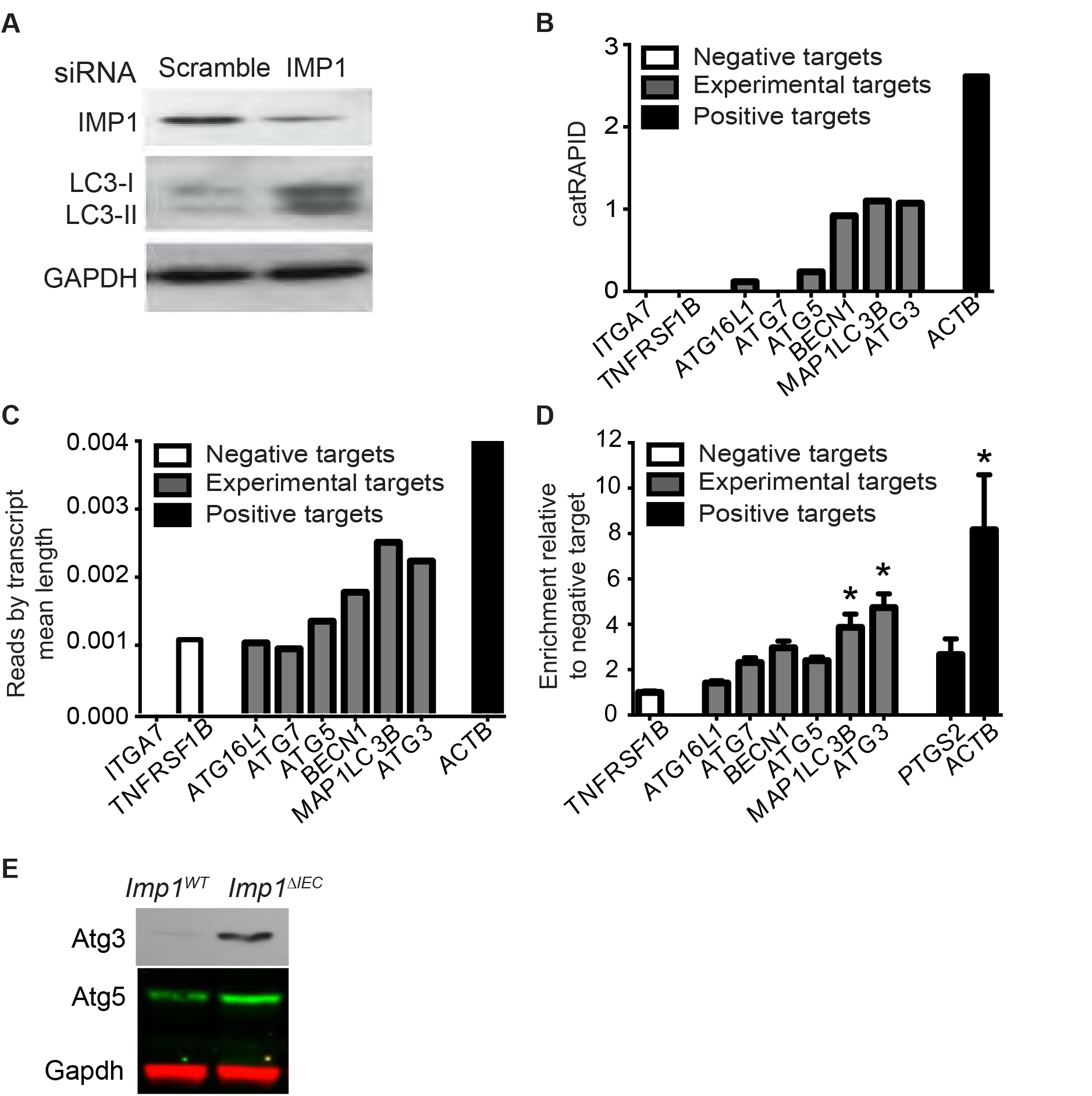
IMP1 interacts with autophagy transcripts. We utilized computational predictions, analyses of published CLIP-Seq data and ribonucleoproteinimmunoprecipitation (RIP) to identify direct interactions between IMP1 and autophagy transcripts. **A**. Caco2 cells transfected with IMP1 siRNA demonstrated a robust increase in LC3-I/LC3-II indicating enhanced flux. Blots are representative of 3 independent experiments. **B**. catRAPID Transcript Score. For each predicted transcript, we measured the IMP1 interaction propensity with respect to the negative control IgG. Negative targets (ITGA7 and TNFRSF1B) were also evaluated. **C**. We retrieved CLIP scores from published eCLIP data against the same set of autophagy-related transcripts analyzed in **B** and found a strong correlation between catRAPID scores and eCLIP data (r =0.9838465 Pearson correlation coefficient). The CLIP data scores were calculated as total number of reads corresponding to the transcript divided by the length of the different isoforms. **D**. We evaluated binding of endogenous IMP1 to autophagy transcripts using RIP assays in Caco2 cells. Specific enrichment of IMP1 was confirmed by IP with either IMP1 or control IgG antibodies followed by western blot for IMP1. Enrichment of target transcripts over control is represented relative to negative target, TNFRSF1B. Positive controls were PTGS2 and ACTB. vs. negative target by 1-way ANOVA. n=3 independent experiments. **E**. Representative western blot showing upregulation of Atg3 and Atg5 in colon epithelium of *Imp1^ΔIEC^* mice as compared to controls. (All data are expressed as mean ± SEM. *, p < 0.05; by ordinary oneway ANOVA test).

To confirm IMP1 binding to these targets, we performed ribonucleoprotein (RNP)-immunoprecipitation with antibodies to endogenous IMP1 in Caco2 cells (Fig. S4A). Previously confirmed IMP1 targets *ACTN* and *PTGS2* and non-target *TNFRSF1B* were used as positive and negative controls, respectively. We observed significant enrichment of IMP1 binding to autophagy genes *MAP1LC3B* and *ATG3* (Fig. 3D). *ATG7, BECN1*, and *ATG5* all demonstrated enriched binding similar to *PTGS2*, which was confirmed as an IMP1 binding target in prior published studies (Manieri et al., 2012). In addition to our initial observation for increased LC3 protein in isolated mouse colon epithelium (Fig. 2D), we observed increased Atg3 and Atg5 in *Imp1^ΔIEC^* colon epithelium (Fig. 3E), suggesting that IMP1 loss directly affects protein levels of key components in the autophagy pathway. We confirmed increased LC3 expression in SW480 cells as well (Fig. S1B). In addition, *in silico* analysis of individual transcripts predicted binding of IMP1 to 5’ UTRs of *MAP1LC3B, ATG3* and *ATG5* (Supplementary Figure 5B-D), which suggests that IMP1 may putatively serve to transiently repress these transcripts. Taken together, we demonstrate via three independent methods that IMP1 binds directly to specific autophagy transcripts required early in the autophagy cascade. Furthermore, we confirm that IMP1 loss enhances autophagy pathway protein expression, which together with *in vivo* phenotypic data suggests IMP1 as a putative posttranscriptional regulator of autophagy in the gut.

### *Imp1^ΔIEC^* mice exhibit enhanced injury recovery, which is reversed with Atg7 deletion

We reported recently that *Imp1^ΔIEC^* mice recover more efficiently following 12Gy whole body irradiation (Chatterji et al, 2018, *in press*). To evaluate the relative contribution of autophagy to the regeneration phenotype in *Imp1^ΔIEC^* mice, we generated *Imp1^ΔIEC^* mice with genetic deletion of autophagy using *Atg7*-floxed alleles (*Imp1^ΔIEC^ Atg7^ΔIEC^*). ATG7 is an essential component of the ATG conjugation system and is critical for early autophagosome formation (Komatsu, Waguri et al., 2005). In addition, prior studies of intestinal epithelial-specific *Atg7* deletion demonstrated loss of autophagic vacuoles via TEM analysis and a phenotype similar to that of intestinal epithelial-specific knockout of the autophagy genes *Atg16L1* or *Atg5* (Adolph, Tomczak et al., 2013, Cadwell, Patel et al., 2009). We challenged *Imp1^ΔIEC^* and *Imp1^ΔIEC^ Atg7^ΔIE^*mice with 12Gy whole body irradiation and evaluated tissue regeneration via quantification of EdU+ regenerative microcolonies (Fig. 4A). *Imp1^ΔIEC^* mice lost less weight and exhibited a significant increase in number of EdU+ microcolonies compared to *Imp1^WT^* mice, consistent with our prior studies (Fig. 4B-D). *Imp1^ΔIEC^ Atg7^ΔIEC^* mice exhibited more weight loss and fewer regenerating microcolonies than *Imp1^ΔIEC^* mice, suggesting that *Atg7* deletion mitigated the beneficial effects of *Imp1* loss in this context (Fig. 4B-D). Analysis of *Atg7^ΔIEC^* mice revealed no difference in weight loss or Day 4 microcolonies compared to wildtype mice (Fig. S5A, B).

**Figure 4.**
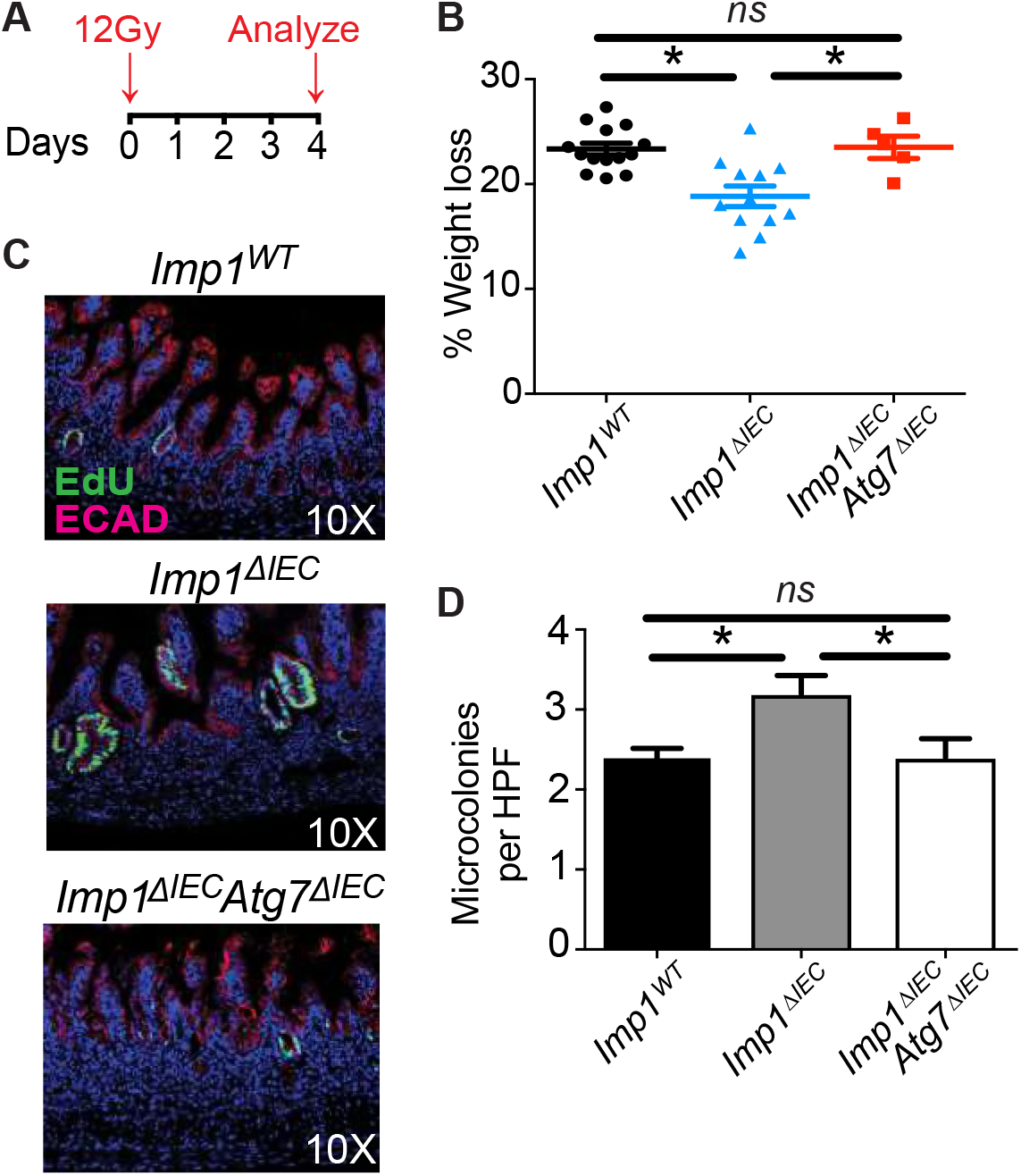
*Imp1^ΔIEC^* mice exhibit enhanced injury recovery, which is reversed with Atg7 deletion. **A**. Simple schematic of the experiment. Mice are treated with 12Gy whole body radiation and all the analyses are performed at day 4 following radiation. **B**. *Imp1* deletion confers protective effects following irradiation, which is reversed in the context of *Atg7* deletion. *Imp1^ΔIEC^* mice lost significantly less weight at sacrifice following irradiation than controls (18.83±0.98% in *Imp1^ΔIEC^* mice(n=12) versus 23.34±0.56% mean weight loss in controls (n=14). This phenotype was abrogated in *Imp1^ΔIEC^ Atg7^ΔIEC^* mice (23.5±1.05% mean weight loss, n=5). For untreated animals, there was no significant difference in mean body weights between groups (not shown). **C&D**. Analysis of EdU+, S-phase cells revealed similar staining in all non-IR mice (not shown); however, there was a robust increase in EdU+ regenerative crypt foci at 4 days following irradiation in *Imp1^ΔIEC^* mice compared to *Imp1^WT^* mice, and this effect was abolished in *Imp1^ΔIEC^ Atg7^ΔIEC^* mice. (n=4 mice per genotype, 20–30 HPF per animal). (All data are expressed as mean ± SEM. *, p < 0.05; by ordinary one-way ANOVA test or standard t-test).

We next evaluated whether *Imp1* loss could promote enhanced recovery in a second damage model, dextran sodium sulfate (DSS)-induced colitis, which leads to colonic epithelial damage (Fig. 5A). Both *Imp1^WT^* and *Imp1^ΔIEC^* mice exhibited significant weight loss following five days of DSS in the drinking water, but *Imp1^ΔIEC^* mice began to recover weight, whereas *Imp1^WT^* mice required sacrifice due to weight loss at Day 9 (Figure 5B). DSS-treated *Imp1^ΔIEC^* mice exhibited significantly lower total colitis and epithelial loss scores compared to controls (Fig. 5C, D, C). Prior studies suggest enhanced susceptibility to colitis in mice with genetic deletion of autophagy (Tsuboi, Nishitani et al., 2015). We observed significant weight loss in both *Atg7^ΔIEC^* mice (Fig. S5) and *Imp1^ΔIEC^ Atg7^ΔIEC^* mice, which became moribund more rapidly than *Imp1^WT^* and *Imp1^ΔIEC^* mice requiring sacrifice prior to the recovery period (Fig. 5B). Thus, while we found that *Imp1* loss lessens acute DSS-colitis, we were unable to ascertain the relative contribution of autophagy to this phenotype. We next evaluated the response of *Imp1^ΔIEC^* mice to chronic colonic epithelial damage (Fig. 5E). *Imp1^ΔIEC^* mice lost significantly less weight following 3 cycles of dextran sodium sulfate (DSS) compared to controls (Fig. S5D). Furthermore, *Imp1^ΔIEC^* mice exhibited an overall decrease in colitis, hyperplasia, inflammation, and mononuclear cell score (Fig. 5F-I). In addition, cytokine expression was decreased in *Imp1^ΔIEC^* mice (Fig. 5J). Together, these observations suggest that *Imp1* loss in intestinal and colonic epithelium promotes more efficient recovery following epithelial injury.

**Figure 5.**
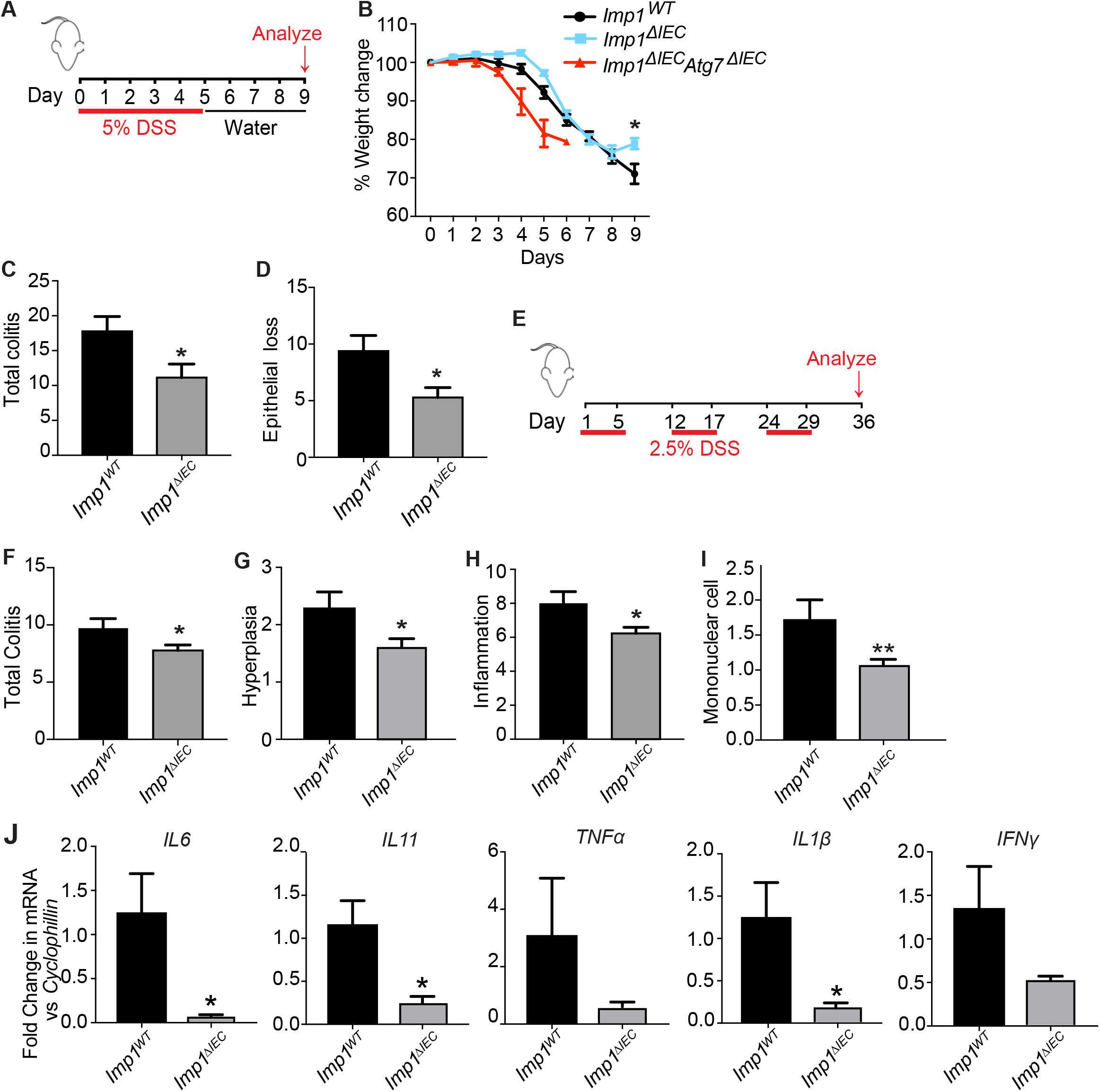
Mice with intestinal epithelial cell deletion of *Imp1* exhibit increased recovery following colitis A. Mice were given 5% DSS in drinking water for 5 days followed by 4 days of recovery. **B**. Mice with *Imp1* deletion lost significantly less weight as compared to controls. *Imp1^ΔIEC^ Atg7^ΔIEC^* mice lost weight and became moribund more rapidly than *Imp1^WT^* and *Imp1^ΔIEC^* mice, requiring sacrifice prior to the recovery period. **C**. *Imp1^ΔIEC^* mice show significantly less total colitis (11.14 ± 1.933, n=7; As scored blinded by a pathologist) as compared to *Imp1^WT^* mice (17.75 ± 2.144, n=8). **D**. *Imp1^ΔIEC^* mice show significantly less epithelial loss (5.286 ± 0.865, n=7) as compared to *Imp1^WT^* mice (9.375 ± 1.375, n=8). **E**. Mice were given 3 cycles of 2.5% DSS in drinking water for 5 days followed by a week of recovery. Mice were analyzed at day 36. **F**. *Imp1^ΔIEC^* mice show significantly less total colitis (7.769 ± 0.4824, n=13; As scored blinded by a pathologist) as compared to *Imp1^WT^* mice (9.714 ± 0.8371, n=7). **G**. *Imp1^ΔIEC^* mice show significantly less hyperplasia (1.615 ± 0.1404, n=13) as compared to *Imp1^WT^* mice (2.286 ± 0.2857, n=7). **H**. *Imp1^ΔIEC^* mice show significantly less inflammation score (6.231 ± 0.3608, n=13) as compared to *Imp1^WT^* mice (8 ± 0.6901, n=7). **I**. *Imp1^ΔIEC^* mice show significantly less mono-nuclear cell infiltration (1.077 ± 0.076, n=13;) as compared to *Imp1^WT^* mice (1.714 ± 0.2857, n=7). **J**. qPCR data showing expression of different cytokines in colon epithelium of *Imp1^WT^* and *Imp1^ΔIEC^* mice at day 36 after chronic DSS treatment. (All data are expressed as mean ± SEM. *, p < 0.05; by standard t-test).

### IMP1 is upregulated in adult and pediatric Crohn’s disease patients

Our data contribute to a working model whereby IMP1 may be induced during stress as a feedback mechanism, exhibiting suppressive effects on stress response mechanisms in gastrointestinal epithelium. We posited that IMP1 expression may be increased as a consequence of chronic stress or disease. We therefore evaluated whole tissue biopsies from adult Crohn’s disease patients, in which we observed a >5-fold increase in *IMP1* compared to control patients (1±0.12 versus 5.3±1.81, Fig. 6A). This was confirmed via IMP1 immunohistochemistry, where we observed both epithelial and stromal IMP1 staining in adult Crohn’s disease patients and confirmed little or absent IMP1 expression in healthy controls (Fig. 6B). Consistent with these findings, analysis of published RISK RNA-sequencing data from pediatric Crohn’s disease (CD) patients revealed that *IMP1* is upregulated significantly compared to control patients, and that this effect is specific to *IMP1* (i.e. other distinct isoforms, *IMP2* and *IMP3*, are not changed; Fig. 6C) (Haberman, Tickle et al., 2014).

**Figure 6:**
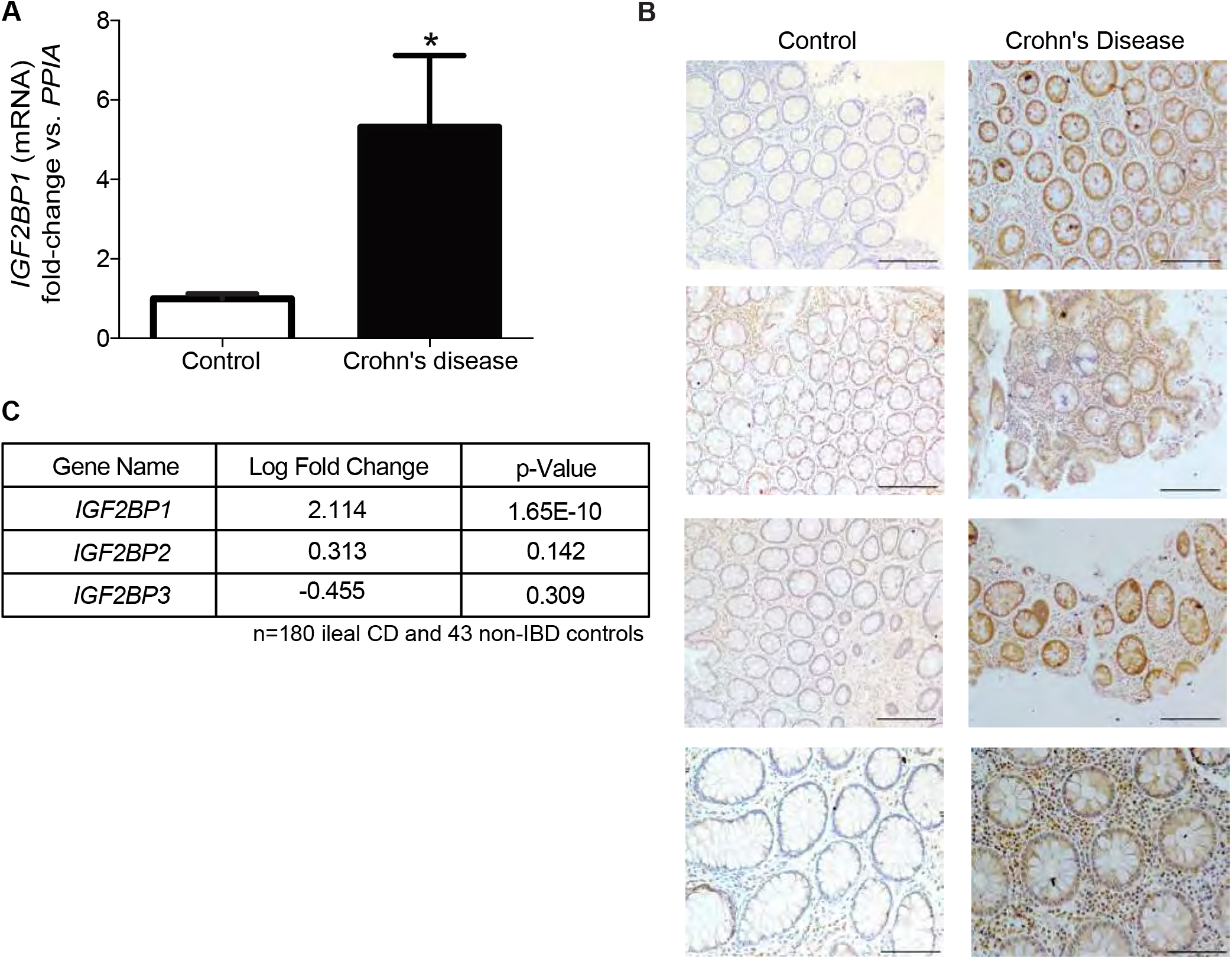
IMP1 is upregulated in adult and pediatric Crohn’s disease patients. **A**.qPCR analysis for *IMP1* expression in colon biopsy samples from adult Crohn’s disease (CD) patients. *IMP1* expression is >5 fold higher in CD samples (5.314 ± 1.807, n=8) as compared to control samples (1 ± 0.1245, n=7). **B**. Representative immunohistochemistry demonstrating IMP1 expression in colon biopsy samples from CD patients and normal adults. IMP1 expression is higher in CD samples (Scale bars = 500m). **C**. Differential gene expression analysis of pediatric CD patient colon samples (n=180) show increased (>4 fold) *IMP1* expression as compared to non-inflammatory bowel disease (IBD) (n=43) pediatric samples. (All data are expressed as mean ± SEM. *, p < 0.05; by standard t-test).

## Discussion

In the current study, we describe a role for the RBP IMP1 to regulate homeostasis in intestinal/colonic epithelium. Unbiased and phenotypic data revealed a mechanistic relationship between IMP1 and the autophagy pathway, underscoring posttranscriptional regulation as an important layer of the gut epithelial stress response. Prior *in vivo* studies have implicated IMP1’s critical role in development, and we and others have demonstrated diverse roles for IMP1 in cancer (Hamilton et al., 2015, Hamilton et al., 2013, Hansen et al., 2004). The present study is the first to uncover *in vivo* mechanisms for IMP1 during intestinal epithelial repair. We demonstrate that *Imp1* loss leads to enhanced repair following damage *in vivo* and that IMP1 is upregulated in Crohn’s disease patients.

IMP1 binds to and regulates the fate of target transcripts via binding to different mRNA regions. Early biochemical studies demonstrated IMP1 binding to the 5′ UTR of the translationally regulated IGF-II leader 3 mRNA, leading to translational repression (Nielsen et al., 1999). IMP1-containing RNP granules are localized around the nucleus and in cellular projections, containing mRNAs representing up to 3% of the transcriptome in HEK293 cells. These granules contain significant enrichment of transcripts encoding proteins involved in ER quality control, Golgi, and secretory vesicles, consistent with our finding that alterations in IMP1 affect the autophagy pathway (Jonson et al., 2007). Furthermore, recent studies describe IMP1 localization in P-bodies, which house mRNAs involved in many regulatory processes, that are transiently repressed (Hubstenberger et al., 2017). This suggests that IMP1 may modulate repression of target mRNAs in certain contexts. Data in the present manuscript suggest that IMP1 may normally serve to repress autophagy and other transcripts.

The levels of functional regulation of autophagy include transcriptional (e.g. TP53, STAT3 and NFκB), posttranscriptional (via miRNAs), and posttranslational (phosphorylation, ubiquitination and acetylation) (Copetti, Bertoli et al., 2009, Feng, Yao et al., 2015, Jing, Han et al., 2015, Lipinski, Hoffman et al., 2010, Yee, Wilkinson et al., 2009). To date, autophagy regulation by RBPs is largely unexplored. Recently, the Dhh1 mRNA decapping regulator and its mammalian homolog DDX6 were implicated as repressors of autophagy through direct modulation of LC3 transcript stability (Hu, McQuiston et al., 2015). This supports the premise that while autophagy activation is beneficial for cellular response to stress, posttranscriptional repressors may play a critical role in attenuating autophagy to prevent prolonged activation. Our biochemical data suggest that IMP1 may represent a new posttranscriptional modulator of the autophagy pathway, acting as an “autophagy rheostat” in intestinal/colonic epithelium.

In IECs, autophagy contributes to microbial handling through packaging and secretion of antimicrobial peptides by Paneth cells. Independent groups have demonstrated that mice with *Atg16l1* gene mutations are more sensitive to colitis or infection, exhibit increased serum IL-1β and IL-18, and display diffuse lysozyme staining in Paneth cells (Burger, Araujo et al., 2018, Cadwell, Liu et al., 2008, Cadwell et al., 2009, Matsuzawa-Ishimoto, Shono et al., 2017, Pott, Kabat et al., 2018, Saitoh, Fujita et al., 2008). In addition, several studies revealed that autophagy genotype-phenotype associations may be used to sub-classify Crohn’s patients (Liu et al., 2016, Rioux, Xavier et al., 2007, VanDussen et al., 2014). Evaluation of *Imp1^ΔIEC^* mice revealed diffuse lysozyme staining. Recent studies have demonstrated that Paneth cells secrete lysozyme via secretory autophagy during bacterial infection through activation of dendritic cell-ILC3 circuit (Bel, Pendse et al., 2017); however, it remains unclear whether secretory autophagy is engaged as a homeostatic mechanism. As such, it would be interesting to determine whether IMP1 may regulate Paneth cell secretory autophagy in future studies.

Finally, our studies demonstrating upregulation of IMP1 in Crohn’s disease suggest increased IMP1 as a consequence of chronic stress or disease. It is yet to be determined if high IMP1 may contribute to Crohn’s disease pathogenesis. Further, the upstream signaling pathways that determine the fate of IMP1-bound transcripts (stabilization versus degradation) within specific contexts, as well as the ability of specific sequence motifs to confer IMP1 target specificity for autophagy (or other) transcripts, remains unknown. Finally, the role for cooperation between IMP1 and microRNAs to elicit specific phenotypes, including effects on autophagy, remain to be determined (Elcheva et al., 2009). In summary, the current study reveals that IMP1 may function as a mediator of epithelial repair in part by modulating autophagy in intestinal epithelial cells. More broadly, these studies underscore the importance of evaluating posttranscriptional contributions to gastrointestinal homeostasis and disease.

## Materials and methods

### Cell lines

Human colon cancer cell line SW480 (ATCC CCL-228) and Caco2 (ATCC HTB-37) cells were obtained from ATCC. Cells are tested for mycoplasma at least every 3 months. *IMP1* was deleted in SW480 cells by cotransfecting cells with IMP1 CRISPR/Cas9 KO Plasmid (h) (Santa Cruz; sc-401703) and IMP-1 HDR Plasmid (h) (Santa Cruz; sc-401703 HDR) followed by sorting and clonal expansion of RFP+ve cells. Knockout was verified by western blotting for IMP1. *IMP1* siRNA (h) (Santa Cruz, sc-40694) or Silencer^®^ Select Negative Control No. 1 siRNA (Thermo Fisher Scientific, 4390843) was transfected in CaCo2 cells using Lipofectamine^®^ RNAiMAX (Thermo Fisher Scientific, 13778075).

### Ribosome profiling

Ribosome profiling libraries from 3 pooled cell culture plates were prepared using a standard protocol (McGlincy & Ingolia, 2017), with minor modifications. Separate 5’ and 3’ linkers were ligated to the RNA-fragment instead of 3’ linker followed by circularization (Subtelny, Eichhorn et al., 2014). 5’ linkers contained 4 random nt unique molecular identifier (UMI) similar to a 5 nt UMI in 3’ linkers. During size-selection, we restricted the footprint lengths to 18–34 nts. Matched RNA-seq libraries were prepared using RNA that was randomly fragmentation by incubating for 15 min at 95C with in 1 mM EDTA, 6 mM Na2CO3, 44 mM NaHCO3, pH 9.3. RNA-seq fragments were restricted to 18–50 nts. Ribosomal rRNA were removed from pooled RNA-seq and footprinting samples using RiboZero (Epicenter MRZH116). cDNA for the pooled library were PCR amplified for 15 cycles. RNA-seq and footprinting reads were mapped to the human transcriptome using the riboviz pipeline (Oana Carja, 2017). Complete TE analyses and pathway analyses are provided in Tables S1 & S2.

### Mice

Mice were cared for in accordance with University Laboratory Animal Resources requirements under an Institutional Animal Care and Use Committee-approved protocol. *VillinCre;Imp1-floxed* (*Imp1^ΔIEC^*) mice were generated previously (Hamilton et al., 2013) and maintained on a C57Bl/6 background. Control mice had floxed, intact alleles (*Imp1^WT^*). Male and female mice were both used at 8–12 weeks. *Atg7-floxed* mice were kindly provided by RIKEN BRC through National Bio-Resource Project of MEXT, Japan (Komatsu et al., 2005). Genotyping primers are listed in Table S3. Mice were housed in specific pathogen-free conditions and fed standard, irradiated chow and water *ad libitum*.

Co-housed control and experimental genotypes were randomized at weaning across multiple cages. For irradiation experiments, animals were given a single dose of 12Gy using Gammacell 40 Cesium 137 Irradiation Unit. Mouse jejunum was analyzed for irradiation experiments. Mice were given 5% dextran sodium sulfate (40,000–50,000 kDa molecular weight; Affymetrix CAS 9011–18-1) in drinking water for acute or 2.5% DSS for chronic colitis (Fig. 5). During all experiments, body weights were recorded daily, and mice were euthanized before losing a maximum of 25% total body weight. Histological scoring was performed blinded by expert veterinary pathologist Enrico Radaelli according to published protocols (Washington, Powell et al., 2013). Sample sizes were determined based upon the investigators’ prior experience with specific models (KEH, GDW). Animal numbers are listed in Table S4.

### Histology

Small intestines were fixed in 10% formalin, processed and paraffin-embedded. Immunofluorescence (IF) staining was performed using heat antigen-retrieval in citric acid buffer (pH 6.0) and staining with antibodies listed in Table S5. 5-ethynyl-2´-deoxyuridine (EdU) staining was performed using Click-iT^®^ EdU Alexa Fluor^®^ 488 Imaging Kit (C10337) as per manufacturer’s protocol. For IMP1 IHC, blocking was performed using Animal-Free Blocker (Vector Laboratories). For all staining, no-primary and/or biological negative controls (*Imp1^ΔIEC^*) were used. Lysozyme scoring was performed according to published protocols (Adolph et al., 2013, Cadwell et al., 2008). Scoring of EdU-positive “microcolonies”, where a microcolony is defined by a cluster of ≥5 EdU-positive cells from a single clone, were quantified, blinded, across at least 30 high-powered fields per animal for a total of 4 mice per genotype.

### Western blot

Caco2 cells or isolated mouse epithelial cells were harvested in western lysis buffer, resolved in reducing conditions on 4% to 12% gradient gels, and detected with ECL Prime Western Blotting Detection Reagent (Amersham; RPN2232). Antibodies are listed in Table S5. Western blots were reproduced in at least three independent experiments.

### Autophagy analyses via CytoID

CytoID Autophagy Detection Kit (Enzo Life Sciences) was used to stain single cell suspensions of crypt-enriched intestinal epithelium (1:100 in DPBS supplemented with 10% FBS at 37°C for 30 min) and co-stained with DAPI. Flow cytometry was performed using FACSCanto or LSR II cytometers (BD Biosciences) and FlowJo software (Tree Star). Unstained cells from each specimen were utilized to establish background fluorescence. The percent of CytoID-positive cells was determined in the live cell fraction (DAPI-negative). The geometric mean fluorescence intensity for live cells was determined for each specimen following subtraction of background fluorescence. Blinded scoring was utilized (KAW) (Merves, Whelan et al., 2016, Whelan, Merves et al., 2017).

### Transmission electron microscopy

Mouse small intestine tissues were fixed in cacodylate-buffered 2.5% (w/v) glutaraldehyde, post-fixed in 2.0% osmium tetroxide, then embedded in epoxy resin and ultrathin sections post-stained in the University of Pennsylvania Electron Microscopy Resource Laboratory. Images were obtained using Jeol-1010 transmission electron microscope fitted with a Hamamatsu digital camera and AMT Advantage imaging software. A total of 4 mice per genotype were evaluated by two investigators for Paneth cell granule morphology and representative photos presented (KEH and BJW) (KlionskyAbdelmohsen et al., 2016). Image contrast was enhanced equally in all TEM photos.

### catRAPID analyses

We used the *cat*RAPID *fragment* approach (Bellucci, Agostini et al., 2011, Cirillo, Agostini et al., 2013) to predict IMP1 binding to autophagy-related transcripts (i.e.: *MAP1LC3B, ATG3, BECN1, ATG5, ATG16L1* and *ATG7*). *ACTB* and *TNFRSF1B/ITGA7* were also included as positive and negative controls, respectively. In our analysis, we included different isoforms for each transcript, for a total of 25 different targets, including the positive and negative controls (http://s.tartaglialab.com/page/catrapid_group).

### Transcript Score

Given a transcript isoform, we used catRAPID *uniform fragmentation* to generate *j* overlapping fragments that cover the entire sequence. The fragments *r_i,j_* are then used to compute *cat*RAPID interacting propensities with IMP1 and IgG (negative control). We define the *interaction threshold θ*(*r_i_*) as the highest interaction propensity score that IgG has with the fragments generated from sequence *r_i_*: 
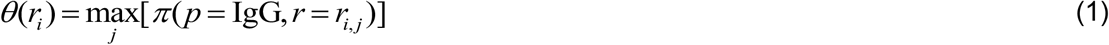

For every IMP1 interaction with fragments, we computed the *normalized interaction π′* by subtracting the interaction threshold of the corresponding transcript to the *cat* RAPID interaction score. 
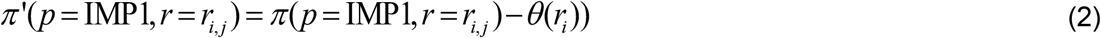

Fragments with normalized interaction score *π′* > 0 are predicted to interact with IMP1. The *Isoform Score* of each isoform ∏*_i_* (Fig. S4*A*) is computed as the average normalized interaction score of interacting fragments *π′* 0over the total number of fragments *n*(*i*): 
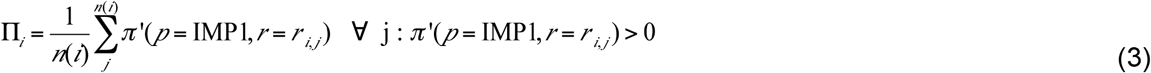

We define the *Transcript Score* ∏_*r*_ (Fig. 5*A*) as a global interaction propensity of all isoforms belonging to a certain transcript. The *Transcript Score* is defined as the average of the Isoform Scores for all the isoforms analyzed for each transcript: 
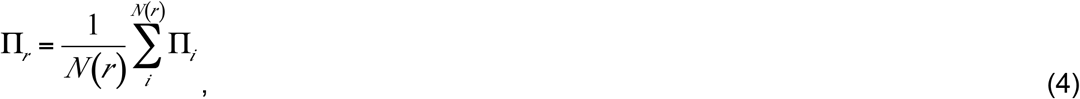
where *N*(*r*) is the number of isoforms considered for each transcript.

### Ribonucleoprotein particle (RNP)-immunoprecipitation

RNP-immunoprecipitations (RIPs) were performed in Caco2 cells using the RiboCluster Prolifer™ RIP-Assay Kit (Medical & Biological Laboratories) according to manufacturer’s instructions. Anti-IMP1 (MBL RN007P, which targets 561–577aa) or control IgG (supplied in kit) were used. Quality control samples for total protein and RNA input as well as immunoprecipitated proteins were evaluated for each experiment. Isolated RNA was reverse-transcribed using the Taqman RT Reagents kit and qPCR performed using the oligonucleotides listed in Table S6. Raw Ct values from IMP1-and IgG-immunoprecipitated samples were used to determine “percent input” for each target, followed by dividing IMP1-immunoprecipitated signal by the respective IgG signal. Data for each individual target were then expressed as fold-enrichment relative to negative control targets *TNFRSF1B*. Positive controls were previously identified IMP1 targets *ACTB and PTGS2*(Manieri et al., 2012, Ross et al., 1997). Three independent RIP experiments were performed.

### qRT-PCR

Small intestine or colon crypt epithelial RNA was isolated using GeneJet RNA purification kit (ThermoFisher). Equal amounts of total RNA were reverse-transcribed using Taqman RT Reagents kit and resulting cDNA used with Power SYBR Green PCR Master Mix (Applied Biosystems/ThermoFisher) or Taqman Fast Universal PCR Master Mix (Applied Biosystems/ThermoFisher) and validated primer sets (Table S6). Non-reverse transcribed samples were used as no RT controls. Gene expression was calculated using *R* = 2^(–ΔΔC_t_)^ method, where changes in *C*_t_ values for the genes of interest were normalized to housekeeping genes. All experiments were replicated in at least 3 independent experiments with technical replicates (duplicates).

### Human samples

Frozen colon tissue and FFPE samples from adult normal and Crohn’s patients were obtained from the Cooperative Human Tissue Network (CHTN) via the University of Pennsylvania Center for Molecular Studies in Digestive and Liver Diseases Molecular Biology and Gene Expression Core. RNA was extracted from frozen tissue using Trizol (Thermo Fisher, Waltham, MA). Publicly available RNA-sequencing data from the RISK cohort of pediatric ileal Crohn’s (Haberman et al., 2014) was evaluated for differential expression of IGF2BP1 (IMP1), IMP2, and IMP3. Sequenced reads were trimmed using Trim Galore! (version 0.4.4), and aligned to the GRCh37 reference genome using STAR, version 2.5.3a. Uniquely mapped reads were quantified by Ensembl gene IDs using featureCounts from Subread version 1.6.0. Lowly or unexpressed genes were removed from the analysis if they showed less than 2 counts per million in less than 5 samples across all conditions. Read counts were transformed with voom and evaluated for differential expression using limma.

### Statistical analyses

Applying unpaired, two-tailed student’s t-tests or 1-way ANOVA, with P <0.05 as statistically significant, determined statistical significance of comparisons between control and experimental conditions unless otherwise noted in the figures legends. For all analyses, unless noted otherwise, data from a minimum of three experiments are presented as mean ± standard error. Sample sizes for individual experiments, including biological and technical replicates, are described in each figure legend, as well as number of experimental replicates.

### Funding

NIH K01DK100485 (KEH), Crohn’s and Colitis Foundation Career Development Award (KEH), NIH R03DK114463 (KEH); Institute for Translational Medicine and Therapeutics of the Perelman School of Medicine at the University of Pennsylvania (KEH); NIH P30DK050306 and its pilot grant program (KEH); startup funds from the Children’s Hospital of Philadelphia Research Institute (KEH); NIH T32CA115299–09 (SFA); NIH R01DK056645 (PC, RM, SFA); NIH K08DK099379 (BJW); European Union Seventh Framework Programme (FP7/2007–2013), through the European Research Council and RIBOMYLOME_309545 (GGT), the Spanish Ministry of Economy and Competitiveness (BFU2014–55054-P and fellowship to FCS), AGAUR (2014 SGR 00685), the Spanish Ministry of Economy and Competitiveness, Centro de Excelencia Severo Ochoa 2013–2017’ (SEV-2012–0208), NIHF32DK107052 (SFA), NIHK01DK103953 (KAW), NIHR03DK118304 (KAW), HHMI Medical Research Fellows Program (ETL), and Fonds de Recherche en Santé du Québec (P-Giroux-27692 and P-Giroux-31601), NIH R01GM103591 (GDW). NIH R35GM124976 (SL, HRSW, PS), NIGMS T32GM008216–29 (SWF), start-up funds from Human Genetics Institute of New Jersey and Rutgers University (SL, HRSW, PS).

## Acknowledgements

We wish to thank Dr. Anil K. Rustgi and lab for support, discussions and technical input. We thank UPenn Core Facilities: Molecular Pathology and Imaging, Human and Microbial Analytic Repository, Cell Culture/iPS, Flow Cytometry, and Electron Microscopy. We thank also Drs. T. Stappenbeck, A. Rodriguez, P. Vedula, L. Ghanem, and Y. Barash for discussions and advice. We thank Drs. Speigelman and Noubissi for *Imp1-floxed* mice. Research reported in this publication was supported by the National Center for Advancing Translational Sciences of the National Institutes of Health under Award Number UL1TR001878. The content is solely the responsibility of the authors and does not necessarily represent the official views of the NIH.

## Author Contributions

Conceptualization: PC, KAW, KEH. Software and Formal Analyses: DSML, SL, SW, FCS, GGT, SM, PS. Investigation: PC, KAW, SFA, FCS, LAS, RM, ETL, SL, SW, SM, LC, PAWVG, BJW, KEH. Writing-Original Draft: PC, KEH. Writing-Review and Editing: PC, KAW, SFA, GGT, PS, GDW, KEH. Funding acquisition: KEH, GGT, PS.

## Conflict of interest

The authors have no conflicts of interest.

